# Multimodal finger-printing of the human precentral cortex forming the motor hand knob

**DOI:** 10.1101/2020.02.11.942771

**Authors:** Raffaele Dubbioso, Kristoffer Hougaard Madsen, Axel Thielscher, Hartwig Roman Siebner

## Abstract

Transcranial magnetic stimulation (TMS) of the precentral hand knob can evoke motor evoked potentials (MEP) in contralateral hand muscles. Biophysical modelling points to the dorsal premotor cortex (PMd) in the superficial crown-lip region as primary site of TMS-induced neuronal excitation. Here, we used a sulcus-aligned MRI-informed TMS mapping approach to determine the optimal site (hotspot) for evoking MEPs in the precentral hand knob. Individual precentral hotspot location varied along the rostro-caudal axis. Individuals with a more rostral location had longer MEP latencies. Spatiotemporal “hotspot rostrality” was associated with higher precentral myelin-related signals, stronger movement-related activation of PMd in the precentral crown, and higher temporal precision during paced finger tapping. Together, our multimodal mapping approach provides first-time evidence for behaviourally relevant, structural and functional phenotypic variation in the crown of human precentral motor hand knob. The results have important implications for physiological and interventional TMS studies targeting the precentral hand knob.

## Introduction

In human and non-human primates, the independent use of single fingers is intimately linked to direct cortico-motoneuronal control of hand and finger muscles (1). The hand representation of the primary motor cortex (M1_HAND_) controls independent fingers movement through fast-conducting, large-diameter pyramidal output neurons that make direct synaptic projections to alpha motoneurons in the lower cervical cord (2). These cortico-motoneuronal axons can be activated non-invasively in awake human subjects using transcranial magnetic stimulation (TMS) (3–6). When TMS is given to the hand representation of the precentral cortex at a sufficiently high intensity, a series of descending volleys are elicited in the corticospinal tract (7). These corticospinal volleys trigger action potentials in spinal motoneurons, producing short-latency motor evoked potentials (MEP) in contralateral hand muscles. Latency measurements of the corticospinal volleys revealed that TMS does not excite the cortico-motoneuronal pyramidal cells directly but via a trans-synaptic indirect mechanism, pointing to a primary site of stimulation that is located “up-stream” to the fast-conducting corticospinal output neurons in M1_HAND_ (6, 7).

The primary site of TMS-induced axonal excitation is largely determined by the absolute magnitude of the TMS-induced electrical field (8). This also applies to the precentral gyrus hosting the motor hand representation. The preferential site of direct TMS-induced excitation is the crown-and-lip region, because the electrical field strength is highest in this most superficial part of the precentral gyrus (8–10). Cytoarchitectonic mapping studies showed that the rostral border of the primary motor cortex is located at the surface of the precentral crown close to the midline, but disappears in the rostral bank of the central sulcus in more lateral parts of the hemisphere (11, 12). This explains why the crown and lip of the precentral hand region is mostly occupied by the caudal dorsal premotor cortex (PMd) which belongs cyto-architectonically to Brodmann area 6 (BA6) (11, 12). The border between rostral M1_HAND_ and PMd is rather gradual without a clear demarcation and the density of giant pyramidal cells in cortical layer V decreases gradually, and this caudal-to-rostral gradient displays considerable interindividual variability (11, 12). The PMd contributes to movement selection and execution and displays a rostro-caudal functional gradient with the caudal PMd being more related to motor execution (13, 14). An executive role of caudal PMd in manual motor control is also reflected by its anatomical connectivity pattern in non-human primates. The caudal PMd is tightly inter-connected with M1_HAND_ and its most caudal part contains scattered large pyramidal cells which send axonal projections to the cervical cord via the pyramidal tract (12, 15). In contrast to caudal PMd in the crown-and lip region, the hand representation of the primary motor cortex (M1_HAND_) is buried in the depth of the sulcal wall (11) and therefore, is unlikely to be a primary site of axonal excitation for TMS (16).

To evoke MEPs, TMS is commonly applied at the scalp site, where TMS elicits the largest MEP amplitude in the target muscle, commonly referred to as “motor hotspot” (6, 17). TMS has been extensively used to study the functional topography of corticomotor representations by applying TMS at different scalp positions using a two-dimensional grid (18–20) and a constant orientation of the coil with respect to the scalp. Several TMS mapping studies have consistently found substantial inter-individual variations in precentral motor hotspot location along the rostro-caudal (anterior-posterior) grid axis of the corticomotor maps (21–26). These inter-individual variations in “hotspot rostrality” raises the possibility that the primary target site of TMS in the precentral gyrus may differ across subjects, when the motor hotspot is used to functionally localize the M1_HAND_. Here, we applied a novel sulcus-aligned TMS mapping approach to precisely localize the motor hotspot of two intrinsic hand muscles along the rostro-caudal axis in the crown of the precentral gyrus (27). We hypothesized that in individuals with a more rostral (anterior) precentral hotspot, TMS elicits premotor cortical excitation more up-stream to M1_HAND_ than in individuals with a more posterior (caudal) precentral hotspot. We therefore expected that individuals with a rostral hotspot should also show longer MEP latencies than individuals with a caudal motor hotspot location in the precentral crown. We additionally performed structural and functional magnetic resonance imaging (MRI) of the precentral gyrus. We hypothesized that inter-individual variations of the precentral motor hotspot location are underpinned by inter-individual structural and functional differences in the precentral gyrus. Specifically, we expected that a more rostral (premotor) hotspot location in the precentral gyrus would be associated with stronger precentral myelination as indicated by structural MRI, larger functional activation of caudal PMd and higher temporal precision during a visuo-motor movement synchronization task.

## Materials and methods

### Participants

Twenty-four healthy young adults (12 women and 12 men) with a mean age of 24 years, ranging from 19 to 34 years participated in this study. Participants were consistently right handed as assessed by the Edinburgh handedness inventory (28). Only individuals with little (<2 years) or no formal music training were included. They had no previous history of neurological or psychiatric disorders and were screened for contraindications to TMS (29). They all gave written informed consent to the experimental procedures. The study complied with the Helsinki declaration on human experimentation and was approved by the Ethical Committee of the Capital Region of Denmark (H-15000551).

### Experimental procedures and data acquisition

Experimental procedures are illustrated in Fig.1 and comprised corticomotor TMS mapping as well as structural and functional magnetic resonance (MRI) of the whole brain at 3 tesla.

### Transcranial magnetic stimulation

We used a sulcus-aligned MRI-informed TMS mapping approach to determine the optimal site (hotspot) for evoking MEPs in the left hand (Fig.1). Single biphasic TMS pulses were applied over multiple sites overlaying the right precentral hand knob. TMS was performed with a cooled MC-B35 figure-of-eight coil connected to a MagPro X100 stimulation device (Magventure, Skovlunde, Denmark). We chose a MC-B35 TMS coil, because this coil is small with an average winding diameter of 35 mm diameter to maximize the focality of TMS. Participants were seated in an adjustable armchair with the neck supported by a head-rest during TMS mapping. The position of the TMS coil relative to the participant’s head was continuously tracked in real time with frameless stereotactic neuronavigation (Localite GmbH, Bonn, Germany). Any changes in the TMS position and orientation relative to the scalp were registered and updated online in a 3D space and displayed to the examiner on a screen. Prior to sulcus-aligned TMS mapping, we located the motor hotspot position of the left FDI muscle by trial-and-error with the handle of the coil angled at 45-degree relative to the midsagittal line. We then determined the resting motor threshold (RMT) of the left FDI muscle, using the Maximum-Likelihood Strategy using Parameter Estimation by Sequential Testing (MLS-PEST) approach (30). The TMS-evoked motor responses were recorded with surface electromyography from relaxed left first FDI and ADM using a belly-tendon montage (Ambu Neuroline 700, Columbia, USA). Trial-wise acquisition of MEPs was controlled by Signal software and EMG data were stored on a computer for later offline analysis (Cambridge Electronic Design, Cambridge, UK). Surface EMG signals were recorded at a sampling rate of 10 kHz 10000 and bandpass filtered (20 Hz – 300 Hz) with an eight channel DC amplifier (1201 micro Mk-II unit, Digitimer, Cambridge Electronic Design, Cambridge, UK).

**Figure 1.**
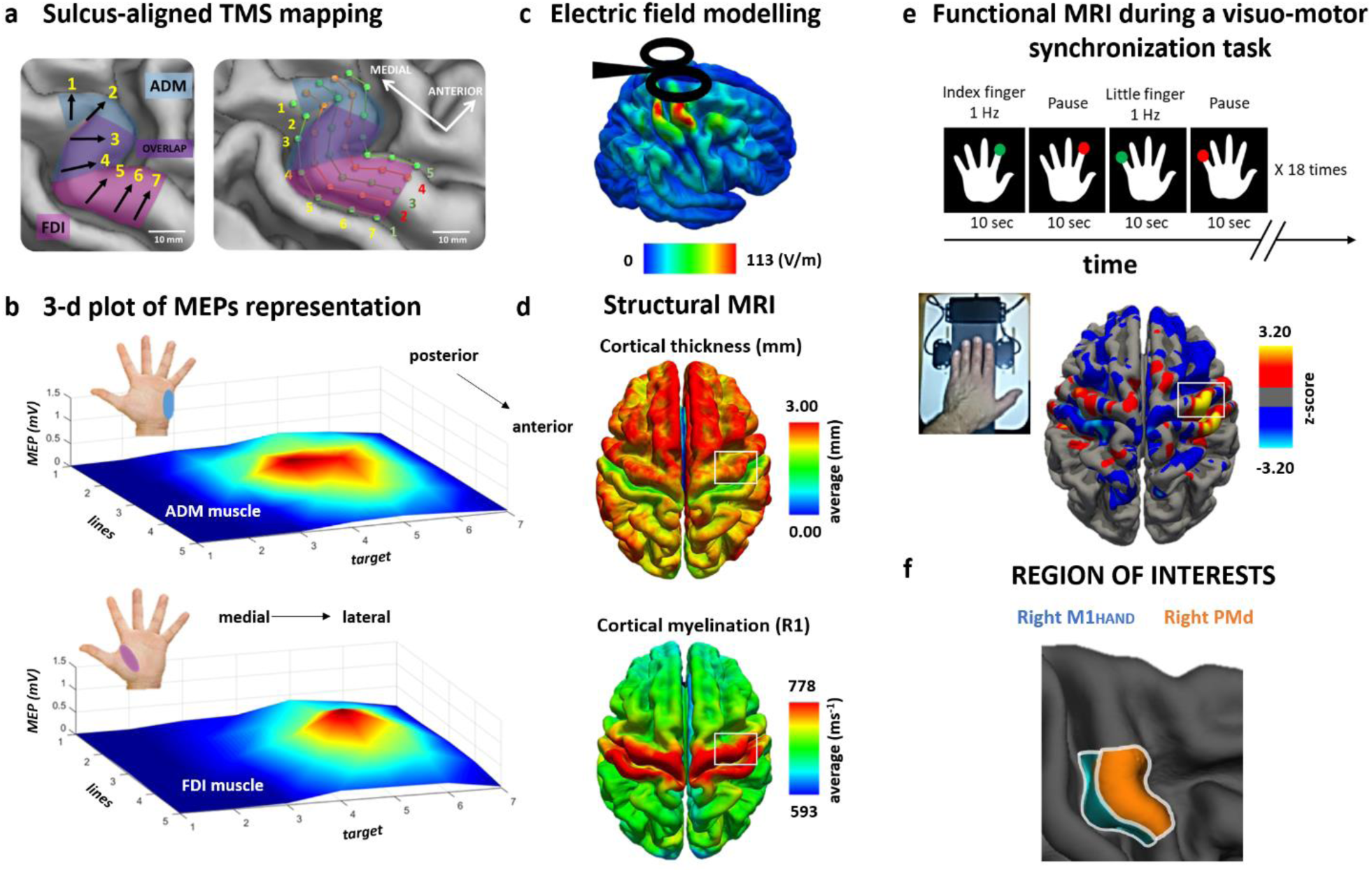
MULTIMODAL PHENOTYPING OF THE PRECENTRAL HAND KNOB. **a** Schematic illustration of sulcus-aligned mapping of single-pulse transcranial magnetic stimulation (TMS). Cortical targets 3, 4 and 5 are located in the centre of the hand knob the macro-anatomical landmark of the M1_HAND_. The blue medial area illustrates the core cortical representation of the abductor digiti minimi (ADM) muscle, while the purple lateral area corresponds to the core cortical presentation of the first dorsal interosseous (FDI) muscle (left figure). Using frameless stereotaxy and based on anatomic landmarks we applied single-pulse TMS to one of the thirty-five cortical target sites (orange and green hexagons) placed along five lines and recorded motor evoked potentials (MEPs) from left first dorsal interosseous (FDI) and abductor digiti minimi (ADM) muscles. The first (most caudal) line corresponded to the border between precentral gyrus and central sulcus. The fifth (most rostral line) corresponded to the border between the precentral sulcus and precentral gyrus. The third line followed the crown of the precentral gyrus. Each target was placed every 5 mm (right figure). Please, note each line follow the central sulcus shape. **b** Three-dimensional surface plot of TMS-induced muscle peak excitability profiles of ADM muscle (top plot) and FDI muscle (bottom plot). Note the FDI muscle is represented more laterally respect to the ADM muscle. **c** Group average of electric field strength |E| at RMT of FDI. **d** Structural MRI: Average distribution of cortical thickness (top) and cortical myelination (bottom) measured as longitudinal relaxation rate R1= 1/T1 across all subjects. **e** Average fMRI activity for voluntary movements of the left index (FDI) and little (ADM) fingers during a visually cued motor task at 1 Hz. **f** Region of interests considered for the analyses: the right primary motor cortex (M1_HAND_) and the right dorsal premotor cortex (PMd) based on the cytoarchitectonic maps.

### Sulcus-aligned TMS mapping of the motor hand knob

Standard grid-based TMS mapping of corticomotor representations keeps the coil orientation of the TMS coil identical across all grid sites (21–26, 31). This procedure induces electrical tissue currents in the motor hand knob that enter the precentral crown at different angles at the maximally stimulated part of the crown, when placing the coil at different points of the grid. This is problematic because the angle at which the electrical current “hits” the precentral gyrus has strong impact on the TMS-induced electric field (32). These considerations prompted us to develop a sulcus-aligned TMS mapping procedure which adjusts the orientation of the TMS coil at each mapping site to the local curvature of the precentral gyrus (27, 33, 34). The procedure exploits the fact that the motor hand representation in the precentral gyrus can be readily identified on structural MRIs by its characteristic knob-like shape (35). The TMS coil is centred on one of seven equidistant target sites placed along the individual gyrus-sulcus border of the hand knob. The coil orientation is adjusted at each target site in a way that TMS always induces the strongest currents in a direction to in the precentral crown that is perpendicular to the local orientation of the precentral gyrus. This secures that TMS induces the highest electrical field strength in the crown of the precentral hand knob at all stimulation sites.

In this study, we extended our linear sulcus-aligned TMS mapping approach into a two-dimensional TMS mapping procedure to identify inter-individual differences in the rostro-caudal peak location of corticomotor excitability in the precentral gyrus. We added four parallel lines rostrally to the central sulcus over the precentral gyrus (Fig.1A). Each of the five parallel lines consisted of seven equidistant targets covering the entire longitudinal extension of the hand knob (35 target sites in total). The distance between two neighbouring target sites on the line was 5 mm. The first (most caudal) line corresponded to the border between precentral gyrus and central sulcus. The fifth (most rostral line) corresponded to the border between the precentral sulcus and precentral gyrus (Fig.1A). The third line followed the crown of the precentral gyrus. TMS mapping was stereo-tactically guided using frameless neuronavigation (Localite GmbH, Bonn, Germany). First, the head of the subject was co-registered with the individual high-resolution anatomical MRI of the brain via anatomical landmarks (e.g., nasion and crus helicis) by using the surface mapping function of Localite. The root mean square differences between positions of landmarks in the MRI volume and at the subjects head were ≤ 2 mm for any TMS session of this study. Furthermore, to verify the quality of the co-registration procedure, we attached small vitamin E capsules (providing a good MRI T1 contrast) to a volunteer’s head at different anatomical positions. The software depicted and true positions of the capsules did not show mismatches larger than 1 mm for any position.

The brain surface from the individual T1-weighted MRI was rendered online and the sites to be targeted by TMS in the precentral gyrus were marked as dots on the segmented brain of each subject (Fig.1A). Based on anatomic landmarks, a trained investigator (R.D.) manually placed thirty-five stimulation sites: seven targets for five lines. As in our recent sulcus-aligned mapping studies (27, 33, 34), we chose a biphasic TMS pulse configuration. Biphasic pulse configurations are more efficient to stimulate the M1_HAND_ than monophasic pulses (36). This allowed to use of a very focal coil without any heating problems, increasing focality compared to standard coils. The second phase of the biphasic stimulus always produced a current in the precentral gyrus with a caudal-to-rostral (posterior-anterior) direction perpendicularly to the local curvature of the central sulcus (37).

To avoid carry-over effects between consecutive stimuli, inter-stimulus intervals were jittered between 4 and 5 s. For each target, we delivered 20 pulses in two 10-stimuli blocks at an intensity set to 120% of the conventionally defined RMT for left FDI muscle. At this stimulation intensity, we reliably evoked motor responses in the left FDI and ADM muscle. The order of stimulated precentral targets was pseudo-randomized across subjects with a fixed target sequence within a subject. The individual coil positioning parameters were stored by the neuronavigation software for each stimulation position (Localite GmbH, Bonn, Germany).

### Magnetic Resonance Imaging

Participants underwent structural and functional MRI the day before the TMS mapping experiment. All MRI scans were acquired with a 3 T Verio Scanner and a 32-channel head coil (Siemens, Erlangen, Germany).

### Structural MRI

Structural T1-weighted images were acquired to assess cortical thickness and to individually identify and track the TMS-target points with frameless stereotactic neuronavigation on each participant’s macrostructure. The T1-weighted images had an isotropic resolution with a voxel size of 1mm^3^ (TR = 2300 ms, TE = 2.96 ms, flip angle = 9°). T2-weighted images were acquired to inform offline simulation of the induced electric fields in the precentral gyrus in each individual participant given the intensity of the stimulation and the distance of the coil from the scalp in each condition. T2-weighted whole-brain scans had a voxel size of 1×1×2 mm^3^ (TR = 10000ms, TE = 52ms, flip angle = 120°). In addition, a whole-brain multiparameter mapping protocol was run to obtain quantitative R1 maps as an index of regional cortical myelination (38, 39). The protocol is based on multi-echo 3D Proton Density- and T1-weighted FLASH (fast low angle shot) images at 1 mm isotropic resolution, which undergo correction for radio frequency (RF) transmit field inhomogeneities using an EPI (echo-planar imaging)-based B1+ map. The latter is corrected for off-resonance effects using a B0 field map. For further details regarding the sequence parameters, we refer to two publications (40, 41).

### Functional MRI

Functional MRI used a gradient-echo planar imaging (EPI) sequence sensitive to detect task-related blood-oxygenation level dependent (BOLD) changes in tissue contrast (TR = 2.07 s, TE = 30 ms, flip angle = 78°, voxel size = 2×2×2 mm^3^, axial field of view = 192×192 mm). A single brain volume consisted of 25 axial slices covering the upper half of the brain. The axial orientation of the brain volumes orientation was slightly tilted backwards so that the orientation of the slices was approximately perpendicular to the course of the central sulcus. 335 brain volumes were acquired during the fMRI session. Two additional short whole-brain EPI scans (62 slices) with same parameters except for an adjusted TR were recorded for co-registration purposes. Pulse and respiration were recorded with an infrared pulse oximeter and a pneumatic thoracic belt. Task related activity changes were mapped with BOLD fMRI while participants performed a repetitive isometric finger abduction task with their left index or little finger (Fig.1E). This task engaged the same muscles that were investigated with TMS, namely the FDI and ADM muscle. The left hand of the subject was placed on a board fitted with two strain gauge sensors measuring the abduction forces produced with the index or little finger (Fig.1E). The strain gauges sensors were connected to custom amplifier which converted the measured force to a voltage in the range 0-2.5 V, this signal was sampled via a PicoLog 1216 USB 2.0 acquisition device at a sampling rate of 500 Hz. Involuntary movement of the thumb, the middle and the ring finger were avoided by using adhesive felt. Subjects saw a schematic drawing of the back of the left hand displayed on a video screen that was visible to the subjects via a coil-mounted mirror. Red or green dots were presented every second at the tip of the left index or little finger. Participants had to produce an isometric abduction with the corresponding finger, whenever a green dot appeared at the tip of the target finger. Participants had to refrain from any movement, whenever a red dot was presented. Using a block design, the same dot and colour was presented 10 times at a constant frequency of 1 Hz and duration of 0.5 s. Blocks of movements (green dots) alternated with blocks without movements (red dots). The four task conditions were always performed in a fixed order and repeated 18 times during the fMRI run (Fig.1E). Stimulus presentation and response recording was controlled by custom-made programs (PsychoPy software, v. 1.74.01, www.psychopy.org) (42). Subjects were familiarized with the task in the scanner, immediately before the fMRI experiment and advised to avoid any additional movements. Performance of the subjects in the scanner was controlled by video monitoring.

## Data analysis

### Motor evoked potentials

After the experiment, the EMG recordings were visually inspected to remove trials with significant artefacts. The peak-to-peak amplitude of each motor evoked potential (MEP) was extracted trial-by-trial using Signal software (Signal Version 4.02 for Windows, Cambridge Electronic Design, Cambridge, UK) in the time window between 10 and 40 ms after the TMS stimulus. For each of the 35 cortical targets, we determined mean peak-to-peak MEP amplitude and used the mean MEP amplitude to generate muscle excitability maps for the ADM and FDI muscles along the precentral gyrus (Fig.1B). The resulting map indicated the site at which the mean MEP amplitude reached its peak. This “MEP peak” indicates the individual location of the motor hotspot in the precentral hand knob. Please note that we used this motor hotspot for further analyses, and not the conventionally identified hotspot location that we had determined by trial-and-error at the start of the experiment to determine RMT. In addition, we tested stability and reproducibility of this procedure on a single subject by replicating the TMS mapping procedure one week later. We used a custom-made software to extract the MNI normalized stereotactic x-, y- and z-coordinates, reflecting the cortical projection of the precentral motor hotspot as revealed by our sulcus-aligned mapping procedure.

We determined the shortest MEP latency for the FDI and ADM muscle for each subject at the motor hotspot location. The shortest latencies were identified and measured by visual inspection of superimposed MEP waveforms (6, 43). This measurement was done by an experienced neurophysiologist (PJS) who was blinded with respect to the protocol set-up.

### Cortical thickness, folding and myelination

Cortical reconstruction was performed with the FreeSurfer image analysis suite ver. 6.0.0 (http://surfer.nmr.mgh.harvard.edu/) (44). Using this approach, the grey and white matter surfaces were defined by an automated brain segmentation process. If required, an experienced investigator manually corrected the automated segmentation, following the procedures outlined at https://surfer.nmr.mgh.harvard.edu/fswiki/Edits. The processes of surface extraction and inflation generated a number of well-known feature descriptors for the geometry of the cortical surface. These included: surface curvature estimated from the mean curvature (or average of the principal curvatures) of the white matter surface (45); cortical thickness estimated at each point across the cortex by calculating the distance between the grey/white matter boundary and the cortical surface. Individual whole brain surface maps were smoothed with a 5 mm 2D Gaussian smoothing kernel (44) and the effect of surface curvature on cortical thickness was regressed out (46). Using the FreeSurfer spherical registration method, the individual curvature-corrected cortical thickness maps were registered to a common FreeSurfer template surface (fsaverage) for visualization and group analysis (Fig.1D) (47).

The data of the multiparameter mapping protocol was processed using a Voxel-Based Quantification (VBQ) toolbox developed of SPM8 (www.fil.ion.ucl.ac.uk/spm/).

Because curvature-associated modulations of myelination can obscure or distort myelination changes due to other variations in cytoarchitectonics (48), we have regressed out curvature and used for analysis the curvature-corrected R1-value variations, smoothed with a 5 mm 2D Gaussian smoothing kernel (44). The individual maps were registered onto the FreeSurfer group template for visualization and group analysis (Fig.1D).

We were primarily interested in estimating cortical thickness in the right precentral gyrus forming the motor hand knob. Therefore, we defined the right motor hand knob as precentral region-of-interest (RoI), covering the M1_HAND_ and the adjacent PMd directly anterior to it. Within this RoI, we separately defined the M1_HAND_ and caudal PMd regions based on the probabilistic cytoarchitectonic maps described in literature for Broadman area 4 and 6 (11, 46, 49). This area was within 2.5 cm of the center of the hand knob (see Fig.1F for visualization). Individual mean cortical thickness values and R1-based cortical myelination estimates were extracted from the two precentral ROIs and used for correlational analyses.

### Task-related fMRI and behavioural data

We used the Statistical Parametric Mapping (SPM) software package (SPM8; Wellcome Department of Imaging Neuroscience, London, UK, http://www.fil.ion.ucl.ac.uk) for pre-processing and statistical analysis of the functional MRI data. The first four volumes of a session (dummy images) were discarded from further analysis. Whole-brain EPIs, including reversed-phase EPI, were recorded to facilitate the accurate registration of the EPI data to the individual T1-weigthed image. The EPI time series was motion corrected, brain extracted and smoothed with a 1 mm two-dimensional Gaussian smoothing kernel (44). We choose a small smoothing kernel to minimize the smearing of task-related somatosensory processing in the postcentral gyrus into the precentral gyrus.

We specified a first-level general linear model to assess differences in brain activity between the movement and rest blocks for each hand muscle. Two regressors-of-interest were defined for the blocks requiring isometric abduction movements of the left index finger (engaging the FDI muscle) or little finger (engaging the ADM muscle). To account for shifts in the onset of the hemodynamic response, temporal derivatives of the resulting time courses, motion, respiration and cardiac cycle were included in the model as regressors-of-no-interest (50, 51). After model estimation, z-statistical images were calculated for the resulting maps of the parameter estimates and a corrected statistical threshold of p<0.05 was applied at the cluster level based on Gaussian random field theory (52). The cluster extent threshold was set to an uncorrected p < 0.001 (here corresponding to a Z-score of 3.2). For reporting, the Z-statistical images were projected into Montreal Neurological Institute (MNI) space based on a nonlinear registration of the T1-weighted structural MRI on the MNI152 template (using FSL FNIRT; http://fsl.fmrib.ox.ac.uk/fsl/slwiki). In addition, average activation maps across subjects were rendered on the FreeSurfer group template (figure 1) for visualization using the registration procedures as described in the online FreeSurfer tutorial (https://surfer.nmr.mgh.harvard.edu/fswiki/FsTutorial/FslFeatFreeSurfer).

During the visually cued isometric finger abduction task, we extracted the mean interval between two consecutive isometric contractions (i.e., inter-movement interval), its standard deviation (SD). Specifically, the signal was thresholded at a level of 0.6 V and the findpeaks matlab function was used to identify peaks with a minimum distance of 400 ms and a peak prominence of 1/3 of the maximum value force value exerted by the subject. As the movement onsets were quite steep, we found it reasonable to use these peaks to define movement intervals. Importantly, the limited dynamic range of the force measurement setup caused the exerted force to often go beyond the maximum, this meant that the force measurements were only of limited practical use in this setting. Lastly, we calculated the coefficient of variation (CV) by dividing the SD with the mean to quantify between-trial dispersion of movement timing. The CV of the inter-movement interval indicates how well participants reproduced the one-per-second interval as signalled by the visual cue.

### Electric field simulations

For each participant, we performed simulations of the electric fields that were generated in the right precentral gyrus by the TMS pulse using SimNIBS software 2.0 (www.simnibs.org). A realistic head model was automatically generated for each participant from the individual T1-weighted and T2-weighted MR images as described elsewhere (53). Electric field simulations were calculated for the coil position which elicited the highest mean MEP amplitude (i.e., the individual precentral motor hotspot) and a stimulation intensity scaled to the individual RMT of FDI to clearly visualize the effect of gyral anatomy on TMS-induced field strength. The vector potential of the MC-B35 coil was pre-calculated using a coil model consisting of a superposition of 1248 magnetic dipoles, as described in Thielscher and Kammer (54). To obtain average electric fields across subjects, the electric fields were interpolated in the middle of the segmented cortical grey matter, and transformed to the FSAverage template (47) for second-level group analyses. We then created a group map of the electrical field distributions for the motor hotspot locations and statistically compared the TMS-induced electrical field distributions in the right precentral hand knob between the M1_HAND_ and PMd group. Since the rostral M1_HAND_ is confined to the posterior lip region of the precentral gyrus (11), we hypothesized that the M1_HAND_ group should display a higher electrical field magnitude in the posterior lip region relative to the PMd group.

### Statistical group analyses

All the statistical computations were performed using IBM SPSS Statistics software (Version 22 for Windows, New York City, USA). Before applying parametric statistical tests, the normal distribution of all variables was verified by means of a Kolmogorov and Smirnov test. Based on the rostro-caudal location of the precentral motor hotspot, we split the population in two groups. Subjects were assigned to the primary motor group (M1_HAND_ group), if the MEP amplitude peak of the sulcus-aligned cortico-motor map of both hand muscles fell in the first two lines close to the central sulcus. Subjects were assigned to the premotor group (PMd group), if the peak location of both muscles was located more rostrally on line three to five. Group demographic data, RMT, shortest MEP latency and MEP amplitude at the motor hotspot were compared between the two groups by using Chi square or unpaired-t tests.

We computed a composite measure that combined the spatial (y-coordinate of hot spot) and temporal dimension (MEP latency) of rostrality. We first normalized each MEP measure (shortest MEP latency and y-coordinate) by scaling it between 0 and 1 and then multiplied them. The composite measure reflects the individual sensitivity of the precentral gyrus to premotor (anterior) or primary motor (posterior) TMS, providing a quantitative spatiotemporal “rostrality index” of precentral excitability. The rostrality index was used to test whether the individual expression of hotspot rostrality is underpinned by structural or functional properties of the precentral gyrus as revealed by structural and functional MRI measurements as well as and task performance during fMRI.

A complementary line of analyses tested for linear correlations across all participants or within the M1_HAND_ and PMd group, calculating the Pearson’s correlation coefficient. Significance threshold was set at p<0.05. Bonferroni procedure was used to correct for multiple comparisons. Data are given as mean (±SEM) if not otherwise specified.

Individual estimates of cortical thickness (derived from T1-weighted MRI scans) and myelination (derived from R1-mapping), mean electric field strengths, and mean task-related BOLD signal increase for index and little finger movements were extracted from the two precentral ROIs in the right precentral hand knob, corresponding to M1_HAND_ and caudal PMd. These variables were analysed in separate mixed-model ANOVAs with the between-subject factor *group* (M1_HAND_ group vs PMd group) and within-subject factors *RoI* (M1_HAND_ vs PMd). *Finger movement* (index vs little finger) was included as additional within-subject factor in the ANOVA models analysing fMRI data and related performance measures. Significance threshold was set at p<0.05.

To visualize the spatial distribution of significant effects within the right precentral hand knob, we computed additional surface-based analyses within the precentral RoI on the FsAverage template by using Freesurfer software. These analyses were performed vertex-wise, followed by cluster-wise corrections for multiple comparisons based on the method suggested by Hagler and colleagues (55) (cluster-determining threshold: p<0.001, clusterwise p<0.05). Age at the time of MRI and sex were included in the model as nuisance variable. Specifically, we used the surface-based analyses to compare the E-field distributions, cortical myelination and functional BOLD activation between the M1_HAND_ and PMd group. Since we had a highly specific anatomical hypothesis (posterior lip region), the explorative between-group analysis of electrical field distribution in precentral gyrus applied a more liberal cluster extend threshold of p<0.01.

## Results

### Precentral corticomotor mapping

Our novel mapping procedure used the so-called “hand knob” as a reliable anatomical landmark on individual T1-weighted MRI brain scans to identify the cortical motor hand representation (35). We stimulated 35 target sites that were located on one of five equidistant parallel lines in the precentral gyrus following the medio-lateral curvature of the precentral motor hand knob. For each target site, the position and orientation of the small focal TMS coil was adjusted with frameless stereotaxy to produce a current direction perpendicular to the curvature of the hand knob at the site of maximal stimulation (27). We calculated the mean peak-to-peak MEP amplitude at each site to create a two-dimensional map of the corticomotor representations of the FDI and ADM muscle (Fig. 2A). This enabled us to identify the location of the precentral motor hotspot for each muscle. The motor hotspot corresponded to the target site, at which TMS evoked on average the largest MEP amplitude. Mean MEP amplitude at motor hotspot was 1.11 (± 0.17) mV for the FDI muscle and 0.50 (± 0.08) mV for the ADM muscle, reflecting the fact that MEP amplitudes were overall larger for the FDI muscle. Extending our previous work, we mapped the spatial representation of MEP amplitudes along five parallel lines in parallel to the curvature of the precentral gyrus. This enabled us to estimate the “rostrality” of the individual hotspot along the anterior-posterior dimension of the precentral gyrus. In agreement with our hypothesis, the position of individual motor hotpots along the rostro-caudal direction varied across participants.

**Figure 2.**
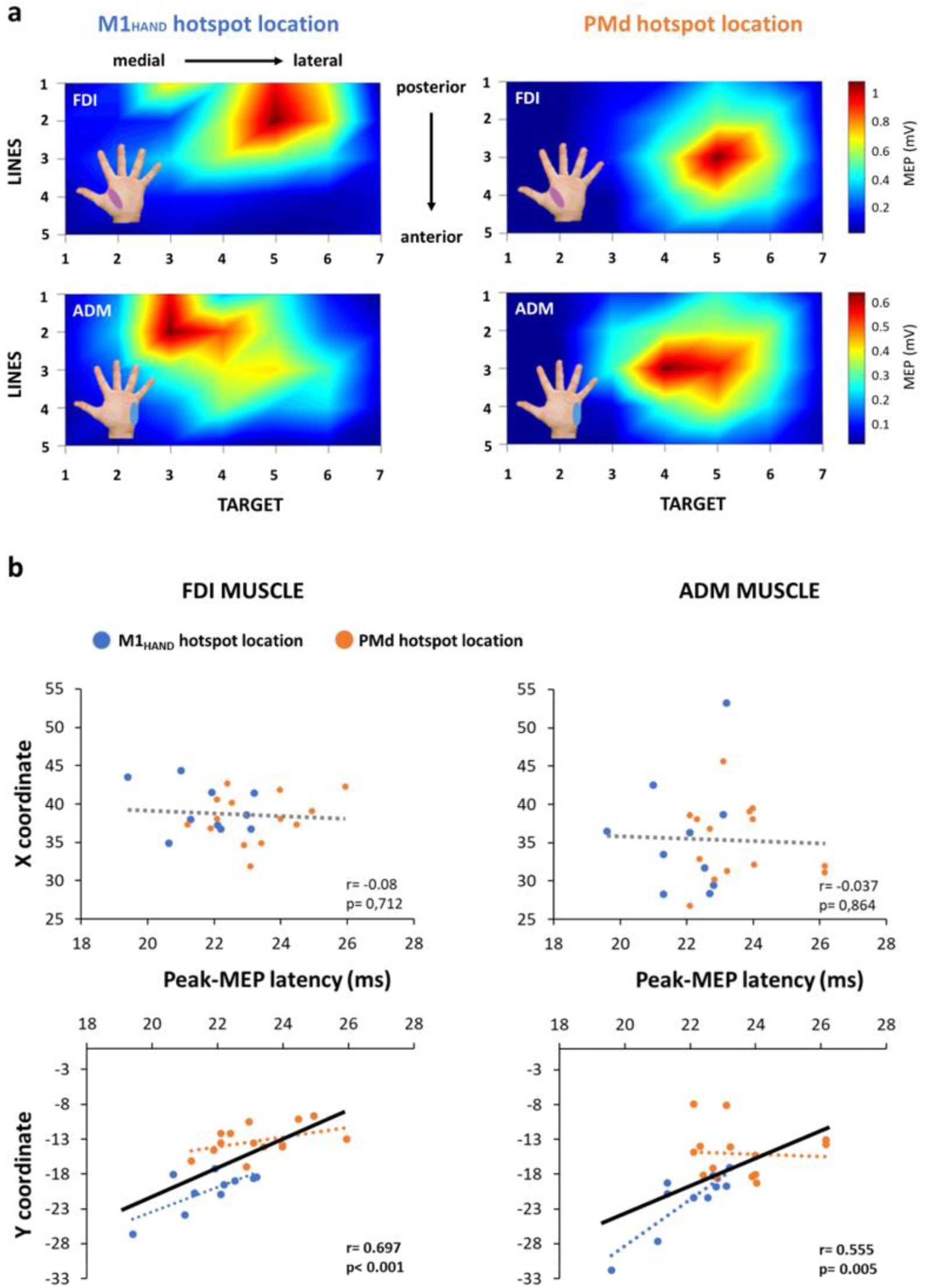
PEAK EXCITABILITY MAPS AND RELATIONSHIP BETWEEN COORDINATES AND MEP LATENCIES. **a** Bidimensional surface plot of TMS-induced muscle peak excitability profiles in subjects with preferential location of hotspot in the primary motor cortex (M1_HAND_) or in the dorsal premotor cortex (PMd) for both muscles (FDI and ADM muscles). Figures indicates the MEPs amplitude profiles (mV) of FDI muscle (top plots) and ADM muscle (bottom plots) along the medio-lateral (target 1 to 7) and posterior-anterior direction (lines 1-5). Note the FDI muscle is represented more laterally respect to the ADM muscle. **b** Linear correlation between X (medio-lateral direction) and Y (posterior-anterior direction) coordinates in the MNI space and latency of the TMS-induced peak excitability recorded from the FDI muscle (left scatter-plot) and ADM muscle (right scatter-plot). Significant correlations with MEPs latencies was found only with the posterior-anterior (Y coordinates) but not with the medio-lateral direction (X coordinates). Blue circles indicate participants with TMS-induced peak excitability in M1_HAND_, orange circles participants with peak excitability in the PMd. Significant correlations are indicated in bold and continuous regression lines (significant p< 0.0125 after Bonferroni correction), non significant correlations by dotted lines.

Based on the rostrality of precentral hotspot location, we assigned participants to a “rostral hotspot” group with the hotspot located on one of the three anterior lines or a “dorsal hotspot” group with the hotspot located on one of the two posterior lines close to the central sulcus. SI appendix, Table S1 summarizes mean group data for the entire group as well as the rostral and caudal hotspot sub-group. Mean MEP amplitudes at hotspot and cortical excitation thresholds for evoking a motor response did not differ in the rostral and caudal hotspot sub-group (p> 0.43), showing that groups were matched in terms of overall efficacy to excite the corticomotor output.

We extracted the stereotactic coordinates to systematically assess the topographical distribution of the motor hotspots in the precentral motor hand knob. Using the stereotactic hotspot coordinates as dependent variable, we computed a mixed-model ANOVA using group assignment as between-subject factor and *hand muscle* (FDI vs ADM muscle) and *axis of stereotactic coordinate* (x-, y-, and z-direction) as within-subject factors. The ANOVA validated our group assignment, showing an interaction between *coordinates* and *group* (F_(2,44)_= 6.049, p=0.005). Post-hoc analyses were performed to test for between-group differences of hotspot locations along the x-, y-, and z-directions. The “rostral hotspot” and “caudal hotspot” groups only differed with respect to their y-coordinates, corresponding to the sagittal (anterior-posterior) dimension in stereotactic space. Mean y-coordinates of both muscles was −21.5 ± 1.3 in the “rostral hotspot” group and −14.1±0.7 in the “caudal hotspot” group (p=0.005). The ANOVA also showed an interaction between *coordinates* and *muscle* (F_(2,44)_= 8.299, p=0.001), confirming a significant difference in precentral location between the FDI and ADM motor hotspots. Specifically, ADM muscle was located more medially (p=0.005) and superiorly (p=0.005) relative to the hotspot of the FDI muscle. This finding replicates our previous sulcus-aligned mapping studies, using a single line of targets placed at the caudal border of the precentral crown (27, 33, 34). Finally, ANOVA also revealed main effects of the two within-subject factors *group* (F_(1,22)_= 47.491, p<0.001) and *coordinates* (F_(2,44)_= 251.644, p< 0.001).

### Spatiotemporal “rostrality” of precentral motor hotspot

To test whether “spatial rostrality” was associated with “temporal rostrality” of the motor hotspot, we tested for a positive linear relation between the antero-posterior coordinate (y) of the motor hotspot and the shortest MEP latency evoked at the hotspot. Individual corticomotor latencies scaled positively with the rostrality of individual motor hotspot locations in the precentral hand knob (Fig.2B). The more anterior (rostral) the motor hotspot was located along the anterior-posterior (sagittal) direction, the longer was corticomotor latency. Considering the data of all participants, we found a significant positive linear relationship between y-coordinate of the motor hotspot and shortest MEP latency at hotspot, both for the FDI muscle (r= 0.697, p< 0.001) and ADM muscle (r= 0.555, p= 0.0045). No such relationship was found for the medio-lateral x-coordinate (Fig.2B) and superior-inferior z-coordinate (FDI muscle: r= 0.084, p= 0.697; ADM muscle: r= 0.109, p=0.614). A significant positive relationship between y-coordinate and MEP latency was also found when only considering the M1_HAND_ group (FDI muscle: r= 0.715, p= 0.02; ADM muscle: r= 0.847, p= 0.002), but not when considering the PMd group (FDI muscle: r= 0.434, p= 0.121; ADM muscle: r= −0.056, p= 0.848). These findings show that “spatial rostrality” is associated with “temporal rostrality”. This association confirms our hypothesis that cortical precentral excitation occurs functionally more “up-stream” to the cortico-motoneuronal neurons, when the preferential target site is located more rostrally in the crown of the precentral hand knob.

Our sulcus-aligned TMS mapping procedure yielded a spatial (y-coordinate) and a temporal (MEP-latency) rostrality measure of the individual TMS target site in the gyral crown of the precentral hand knob. Considering both, the spatial and temporal rostrality dimensions, we computed a “spatiotemporal rostrality index” of the TMS hotspot (see Methods section for details). This spatiotemporal rostrality index reflects how much TMS preferentially excited cortical neurons in the caudal PMd or M1_HAND_. At the individual level, the spatiotemporal rostrality index of the FDI and ADM muscle showed a positive linear relationship (r= 0.819, p<0.001), showing that hotspot rostrality of the two hand muscles was consistently expressed at the single-subject level.

### Rostrality of precentral motor hotspot has a functional correlate

We hypothesized that the rostrality of the precentral motor hotspot would be relevant to manual motor control. We therefore tested whether inter-individual differences in hotspot rostrality would be reflected by differences in task-related regional activation and task performance. Our task-related fMRI measurements yielded evidence in support of both predictions (Fig.4). When performing paced finger movements at a repetition rate of 1Hz, individuals with a rostral motor hotspot activated the PMd more strongly than individuals with a posterior hotspot location in rostral M1_HAND_ (Fig.4A). Mixed-model ANOVA yielded a significant interaction between *Region of Interest* (*RoI)* and *group* (F_(1,22)_= 5.431, p= 0.029). Post-hoc between-group comparison revealed that the PMd group showed a relatively stronger task-related BOLD increase in the PMd RoI relative to the M1_HAND_ group (p= 0.002). The activation level in M1_HAND_ did not differ between the M1_HAND_ and PMd group (Fig.4A). The precise localization of the difference between the two groups was visualized based on a surface-based voxel-wise analysis (SI Appendix, Fig. S1C-D). Surface-based analysis revealed a cluster in the crown of the precentral gyrus with a regional peak at x, y, z = 34.4, −15.4, 65.3 (SI Appendix, Fig. S1C). Considering the data of all participants, we found a significant positive linear relationship between the task-related BOLD increase in the PMd region and the spatiotemporal rostrality index of the precentral motor hotspot (Fig.4B). This was the case for movements with the index finger (r= 0.508, p= 0.011) and little finger (r= 0.611, p= 0.0015). When considering each group separately, the PMd group showed a positive relation between task-related BOLD increase during index finger movement and the rostrality index (r= 0.484, p= 0.017).

**Figure 3.**
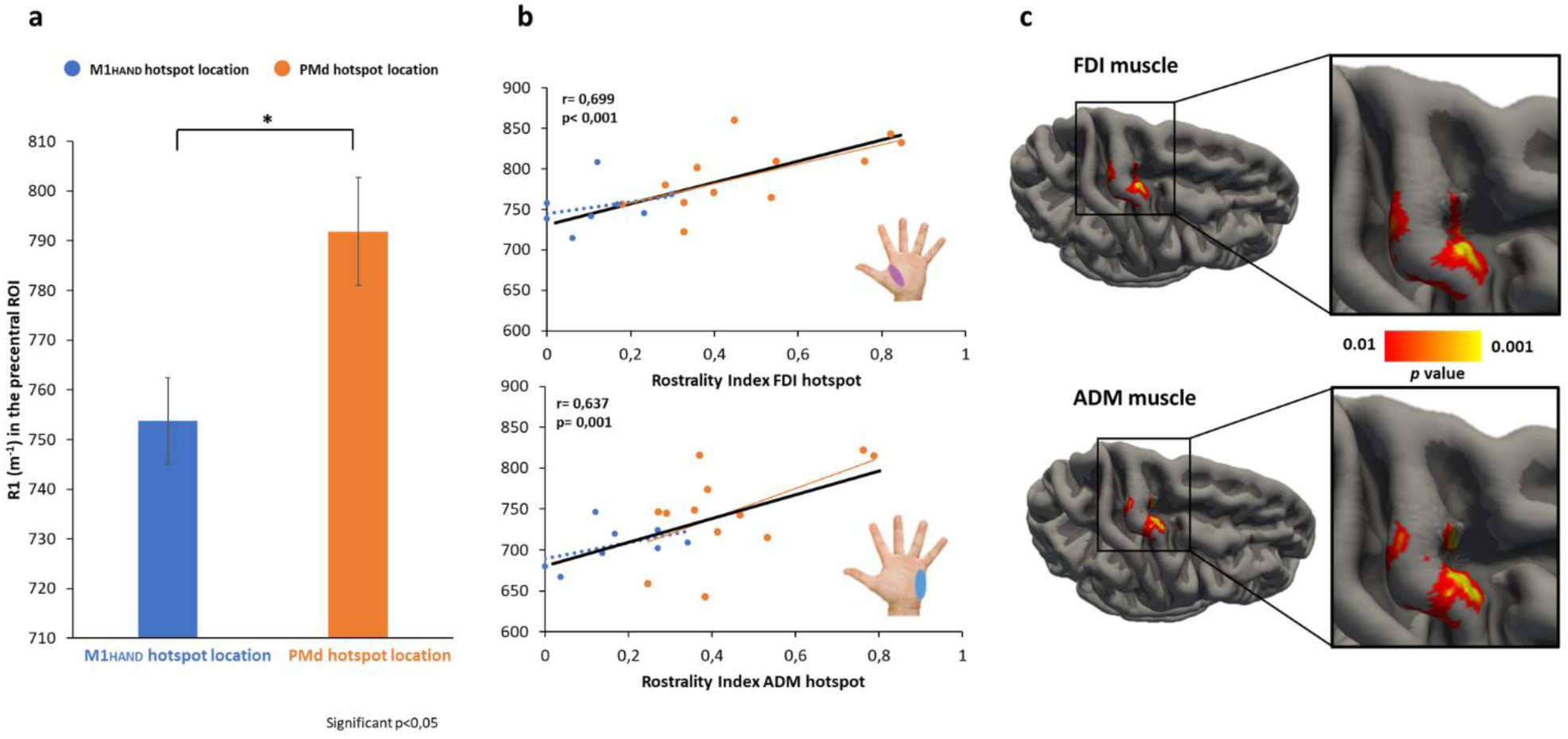
MICROSTRUCTURAL CORRELATES OF THE SPATIOTEMPORAL “ROSTRALITY” INDEX. **a** Between-group difference of R1 estimated cortical myelination in the primary motor (M1_HAND_) and dorsal premotor cortex (PMd) region of interests (Fig. 1F). The mean value of R1 in subjects with preferential hotspot location in the PMd (orange bar) was significantly higher than subjects with hotspot location in the M1_HAND_ (blue bar) (p =0.033). *= statistically significant. **b** Significant linear correlation between R1 estimated cortical myelination and the spatiotemporal “rostrality” of the FDI muscle hotspot (top scatter plot) and ADM muscle hotspot (bottom scatter plot). Significant correlations are indicated in bold and a continuous regression line, non-significant correlations by dotted lines. **c** Surface correlation maps indicating a significant relationship between the spatiotemporal “rostrality” index and cortical myelination for the FDI muscle (top figure) and ADM muscle (bottom figure) (thresholded at p_uncorr_<0.01). Cluster-wise correction showed two main significant clusters in the precentral gyrus for the FDI muscle (Cluster-wise probability, CWP-p= 0.0001) and for the ADM muscle (CWP-p= 0.0001).

**Figure 4.**
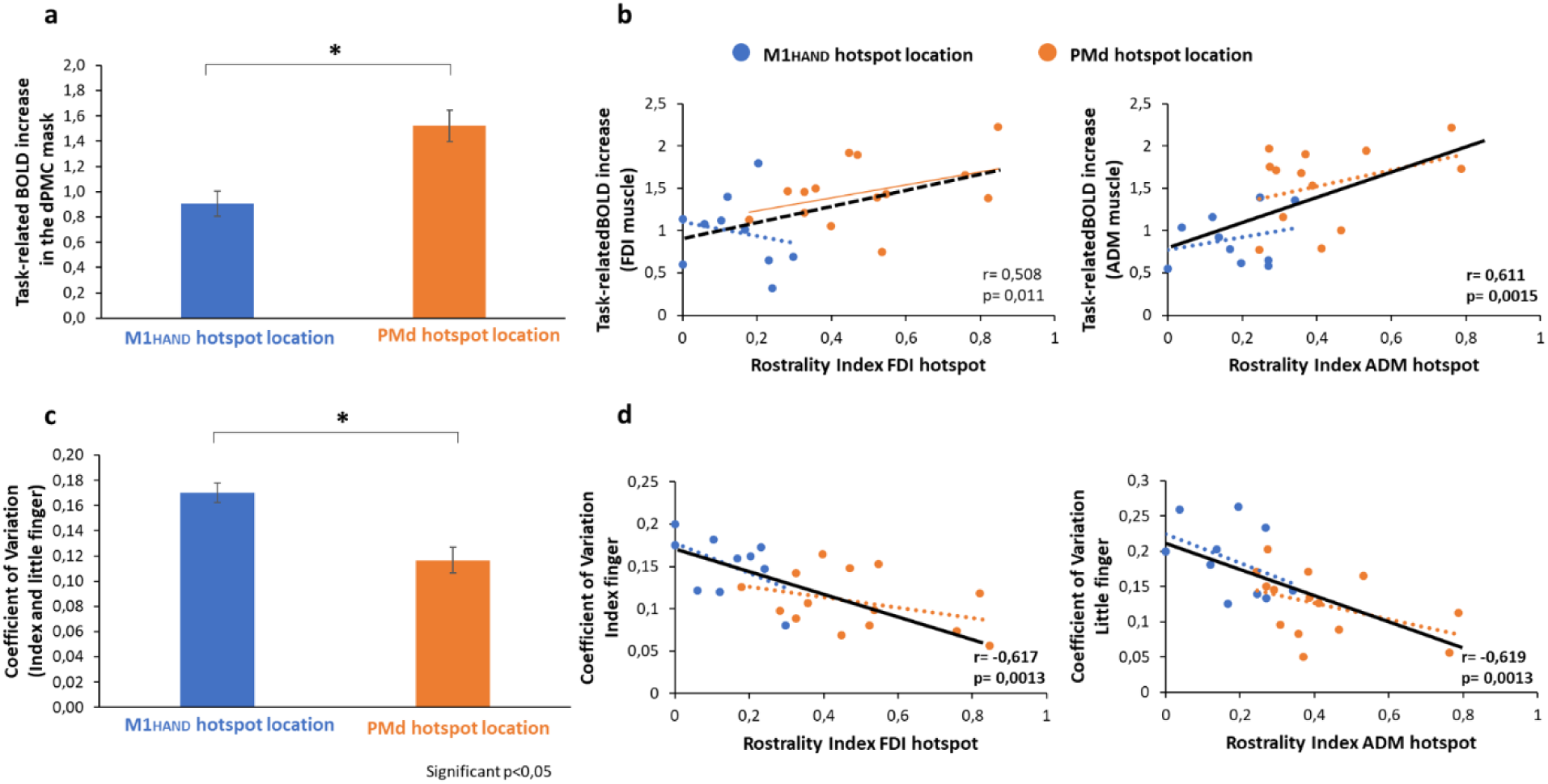
FUNCTIONAL AND BEHAVIORAL CORRELATES OF THE SPATIOTEMPORAL “ROSTRALITY” INDEX. **a** Between-group difference of mean BOLD-increase during a visuomotor-synchronization task at 1 Hz. Individuals with a rostral motor hotspot (orange bar) activated the PMd more strongly than individuals with a posterior hotspot location in rostral M1_HAND_ (blue bar) (post-hoc test of the significant *group* by *RoI* interaction, p =0.002). **b** Significant linear correlation between cortical activation during the visuo-motor synchronization task and the spatiotemporal “rostrality” of the ADM muscle hotspot (right scatter-plot) but not for the FDI muscle hotpsot (left scatter-plot). **c** Between-group difference of coefficient of variation (CV) of the inter-tap interval. The mean CV for both muscles (FDI and ADM) in subjects with preferential hotspot location in the PMd (orange bar) is significantly higher than subjects with hotspot location in the M1_HAND_ (blue bar) (p <0.001). **d** Significant linear correlation between the CV of the inter-tap interval and the spatiotemporal rostrality index for the FDI muscle (left scatter-plot) and ADM muscle (right scatter-plot). Significant correlations are indicated in bold and by continuous regression line, non significant correlation by dotted lines. Black line indicates all participants. *= statistically significant.

Mixed-model ANOVA also revealed a main effect of *finger* (F_(1,22)_= 9.612, p= 0.005) which was caused by a higher task-related BOLD increase during movements with the index finger as opposed to movements with the little finger. The stronger activation during index finger movements was more evident in M1_HAND_ than PMd region (Interaction between *RoI* and *finger*: F_(1,22)_= 13.035, p= 0.002). A significant main effect of *RoI* (F_(1,22)_= 16.411, p= 0.001) reflected a stronger task-related activation of the M1_HAND_ relative to the PMd RoI. Rostrality of precentral motor hotspot also scaled with the regularity of repetitive movements as reflected by the temporal variability of the paced finger tapping movements. The PMd group showed less temporal dispersion of inter-movement intervals, as indexed by a lower coefficient of variation (CV), than the M1_HAND_ group for tapping movements with the index and little finger (Fig. 4C). Accordingly, mixed-model ANOVA showed a main effect of *group* (F_(1,22)_= 15.259, p< 0.001) but no interaction between *group* and *finger (F*_*(1,22)*_*=0*.*841; p= 0*.*369)*. There was also a main effect of *finger* (F_(1,22)_= 5.899, p= 0.024) with the little finger producing more variable inter-movement intervals than the index finger. The temporal variability of the inter-tap interval also displayed a significant negative correlation with the spatiotemporal rostrality index for both fingers across all participants (index finger: r= −0.617, p=0.0013; little finger: r= −0.619, p= 0.0013). This negative linear relationship indicates that participants with lower temporal variability of tapping rate had a more rostral motor hotspot in the precentral gyrus (Fig.4D). This correlation did not reach significance when considering the M1_HAND_ (index finger: r= −0.513, p=0.13; little finger: r= −0.429, p= 0.22) or PMd group separately (index finger: r= −0.365, p=0.2; little finger: r= −0.424, p= 0.13).

### Rostrality of precentral motor hotspot has a microstructural correlate

The structural properties of the right precentral hand knob were assessed with quantitative structural MRI, using the mean R1-value of caudal PMd and M1_HAND_ as index of regional myelination (see methods section for details). The R1-value was only available in 20 of the 24 subjects, as the quantitative MRI data of four subjects had to be excluded because of head movement related artefacts. The PMd group had higher mean R1-values in the precentral cortex than the M1_HAND_ group (Fig. 3A). This between-group difference was reflected by a main effect of *group* in the mixed-model ANOVA (F_(1,18)_= 5.36, p= 0.033). ANOVA also showed a main effect of *RoI* (F_(1,18)_= 24.164, p< 0.001), which was due to stronger cortical myelination of M1_HAND_ (796 ± 8 ms^-1^) relative to PMd (757 ± 9 ms^-1^) region. We computed a surface-based voxel-wise analysis to visualize the regional expression of between-group differences in R1-values. At the voxel level, between-group difference in R1-value was maximal in the anterior lip of the precentral gyrus at the transition zone from the crown of the precentral gyrus to the precentral sulcus (peak difference at x, y, z = 22, −12, 58, see SI Appendix, Fig. S1A-B).

The individual spatiotemporal rostrality index showed a significant positive linear relationship with the degree of precentral myelination as indexed by mean R1-value of the M1_HAND_ and PMd region. The more rostral the precentral motor hotspot, the more myelinated was the precentral cortex (Fig.3B). This positive relationship was found for the spatiotemporal rostrality index of the FDI hotspot (r= 0.699, p<0.001) and ADM hotspot (r= 0.637, p= 0.0019). When considering each sub-group separately, this linear relationship was still present in the PMd group for the FDI hotspot (r= 0.647, p= 0.011) and ADM hotspot (r= 0.564, p= 0.028), but not in the M1_HAND_ group. Surface-based correlation analyses revealed that the local R1-values in a rostro-lateral part of PMd at the anterior lip of the precentral gyrus correlated most strongly with the individual spatiotemporal rostrality index (Fig. 3C; peak correlation at x, y, z = 32.6, −11.3, 53.6 for FDI muscle and x, y, z = 34.8, −11.6, 59.2 for ADM muscle). A second cluster was located dorso-medially in the posterior lip region of the precentral gyrus, (peak correlation at x, y, z = 26.6, −22.7, 51.8 for FDI muscle and x, y, z = 29.3, −21.8, 58.7 for ADM muscle), corresponding to the rostral M1_HAND._

To further explore the impact of precentral myelination on the performance during the visuomotor synchronization task, we performed exploratory correlation analyses within the precentral RoI, including the M1_HAND_ and PMd regions. We found a significant negative relation between precentral cortical myelination (as indexed by regional R1-value) and the mean CV of tapping rate for both fingers in two precentral clusters (Fig. 5A). A first ventrolateral cluster peaked at x, y, z = 38, −16, 57 mm (cluster-wise p< 0.001) and a second dorsomedial cluster peaked at x, y, z = 30, −19, 68 mm (cluster-wise p< 0.001), fig. 5B-C. Both groups contributed to this correlation with r-values above −0.7 at both regional peaks (Fig. 5B-C).

**Figure 5.**
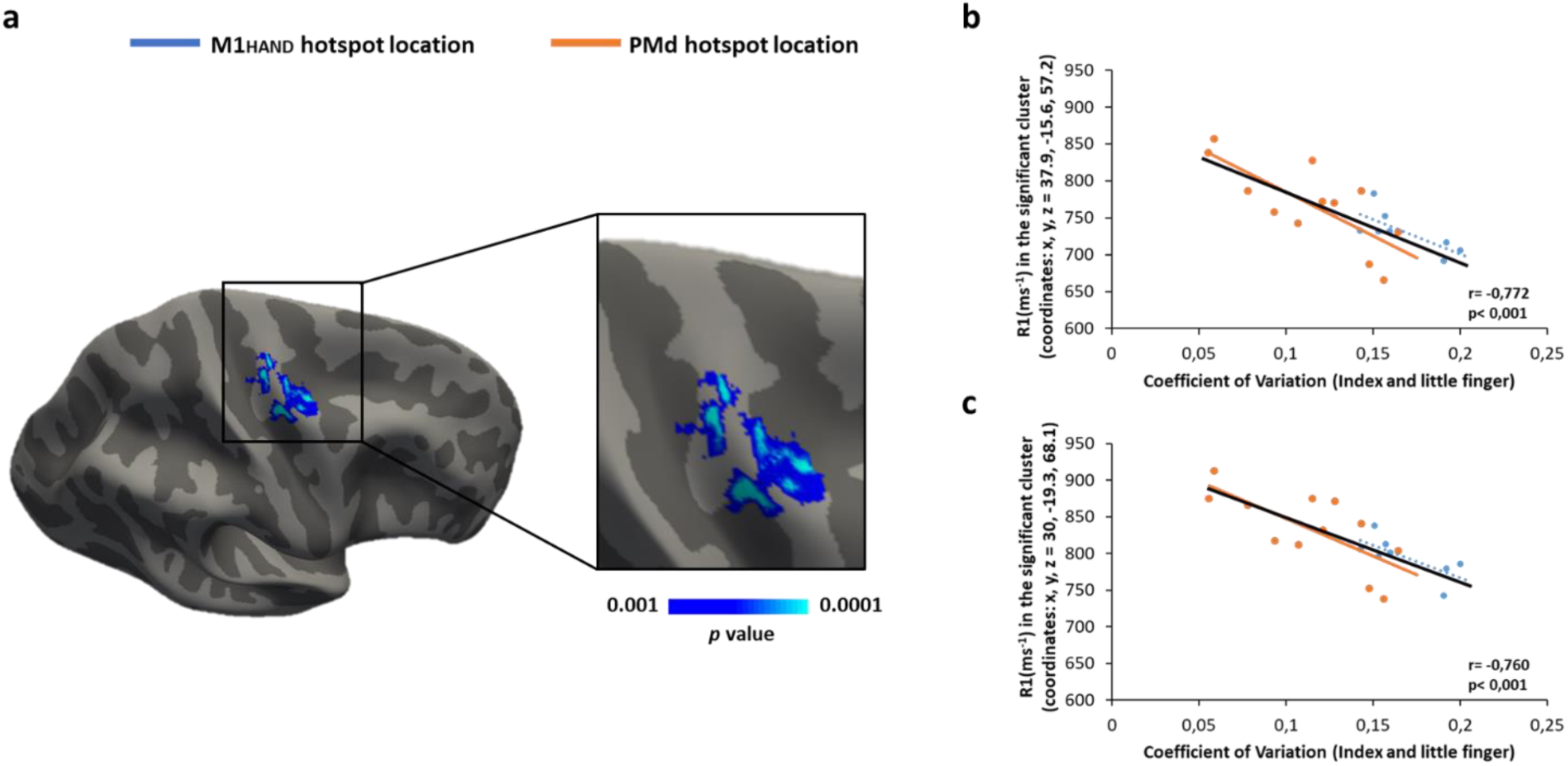
RELATIONSHIP BETWEEN CORTICAL MYELINATION AND HAND MOTOR SKILL. **a** Surface correlation map indicating significant negative relationship between cortical myelination and the mean coefficient of variation (CV) of the inter-tap interval for both fingers (Index and little fingers) within the precentral *RoI* (thresholded at p_uncorr_<0.001). **b** and **c** Significant linear negative correlations between the mean CV of the inter-tap interval and cortical myelination in first ventrolateral cluster (coordinates of most significant vertices: x, y, z = 37.9, −15.6, 57.2) and in the second dorsomedial cluster (coordinates of most significant vertices: x, y, z = 30, −19.3, 68.1) identified by cluster-wise correction for multiple comparisons. Significant correlations are indicated in bold and by continuous regression line, non-significant correlation by dotted lines (p< 0.008 after Bonferroni correction). Black line indicates all participants

In contrast to mean R1-values, regional cortical thickness of right precentral gyrus did not differ between the M1_HAND_ and PMd group. The caudal PMd region was thicker (2.73 ± 0.03 mm) than the M1_HAND_ region (2.51 ± 0.04 mm), but this difference in cortical thickness was comparable between groups. Accordingly, the mixed-model ANOVA with the within-subject factor *RoIs* and between-subject factor *group* only revealed a main effect of RoI (F_(1,22)_= 40.6, p< 0.001), but no significant effect of *group* (F_(1,22)_= 0.137, p= 0.715) or an interaction between *RoI* and *group* (F_(1,22)_= 0.434, p= 0.517). At the individual level, regional thickness of the right precentral gyrus did not predict spatiotemporal rostrality of the precentral motor hotspot. No significant correlation was found between cortical thickness and the individual rostrality index for both muscles (FDI muscle: r= −0.230, p= 0.280; ADM muscle: r= −0.277, p= 0.190).

Cortical folding, indexed by regional mean curvature, of the right precentral gyrus did not differ between the M1_HAND_ and PMd group. As expected, the caudal PMd region exhibited a lower value (−0.08 ± 0.01 mm^-1^) than the M1_HAND_ region (0.02 ± 0.01 mm^-1^), but the difference in cortical curvature was comparable between groups. Indeed, the mixed-model ANOVA with the within-subject factor *RoIs* and between-subject factor *group* only revealed a main effect of RoI (F_(1,22)_= 1709.9, p< 0.001), but no significant effect of *group* (F_(1,22)_= 0.183, p= 0.673) or an interaction between *RoI* and *group* (F_(1,22)_= 0.418, p= 0.525). At the individual level, no significant correlation was found between mean curvature of the right precentral gyrus and the rostrality index for both muscles (FDI muscle: r= 0.146, p= 0.495; ADM muscle: r= −0.021, p= 0.924).

Together, these negative results show that the association between regional myelination and spatiotemporal rostrality of the precentral hotspot was not driven by differences in cortical volume and mean curvature.

### Hotspot location is related to induced electrical field magnitude in precentral gyrus

Since all participants underwent structural T1-weighted and T2-weighted MRI scans, we were able to simulate the electric fields that were generated in the right precentral gyrus by the TMS pulse at the individual precentral hotspot. We created a group map of the electrical field distributions for the motor hotspot locations and statistically compared the TMS-induced electrical field distributions in the right precentral hand knob between the M1_HAND_ and PMd group. Since the rostral M1_HAND_ is confined to the posterior lip region of the precentral gyrus (11), we hypothesized that the M1_HAND_ group would display a higher electrical field magnitude in the posterior lip region relative to the PMd group. Confirming our hypothesis, between-group comparison of TMS-induced electrical field distributions in the right precentral hand knob showed higher electrical field magnitudes in the posterior lip region for hotspot stimulation in the M1_HAND_ group compared to the PMd group (SI Appendix, Fig. S2). The between-group difference in electrical field magnitude peaked at the x-,y-,z-coordinates 34, −20, 65, corresponding to the rostral part of M1_HAND_.

## Discussion

Sulcus-aligned TMS mapping yielded novel insights into the functional neuroanatomy of the human precentral hand knob. We found that the optimal location for precentral TMS to evoke MEPs varies across individuals along the rostro-caudal axis of the precentral hand knob. A more rostral hotspot location in the precentral crown was associated with a longer corticomotor MEP latency. Inter-individual differences in “hotspot rostrality” scaled with structural and functional properties of the precentral gyrus. Hotspot rostrality also had a behavioural correlate. Individuals with a more rostral hotspot displayed higher temporal precision during visually paced finger tapping. Together, our results link a spatiotemporal feature of corticospinal excitability to the structure and function of the precentral cortex. In the following, we first discuss the corticomotor mapping results yielded by sulcus-aligned MRI-informed TMS mapping. We then comment on the potential functional significance of a more rostral motor hotspot for dexterity. We finally highlight the implications of our results for the use of TMS to study and modulate the corticomotor system.

Sulcus-aligned TMS mapping replicated previous grid-based TMS mapping studies, showing that the spatial location of the optimal site for evoking MEPs in the precentral hand knob, the motor hotspot, varied across individuals along an anterior-posterior axis (21–26). Our sulcus-aligned mapping procedure secured that TMS at each stimulation site produced always a consistent current direction in the most strongly stimulated part of the precentral hand knob regardless of the individual folding pattern. The comparable stimulation conditions for all stimulation sites minimized the influence of the curvature of the precentral hand knob on the mapping results. Hence, a more rostral motor hotspot location indicates that the corticomotor output is most effectively excited by TMS in a more rostral site in the precentral crown-and-lip region.

The inter-individual variation in precentral hotspot location has important implications regarding the primary target site of TMS. While the precentral crown-and-lip region is mainly occupied by the PMd (BA6), a part of the posterior lip might belong to the rostral part of anterior M1_HAND_ (BA4a) (11). Hence, TMS with a caudal hotspot in the posterior lip region may primarily stimulate the rostral part of the anterior M1_HAND_ (BA4a), whereas TMS in individuals with a rostral hotspot in the precentral crown may primarily stimulate PMd (BA6). This view is supported by the observation that the group with a caudal hotspot location (referred to as M1_HAND_ group) had higher electrical field magnitudes in the posterior lip region than individuals with a rostral hotspot location.

In this context, it is important to point out that retrograde tracing studies in the macaque monkey identified the caudal portion of M1 in the anterior bank of the central sulcus as the main precentral source of cortico-motoneuronal pyramidal output neurons (56, 57). The rostral portion of M1 in the crown of the precentral gyrus contained only few cortico-motoneuronal pyramidal output neurons (56, 57), indicating a sub-division of the M1 in a rostral “old” and caudal “new” M1 (56, 57). Under the assumption that this sub-division also applies to the human M1_HAND_, it is unlikely that TMS causes significant direct stimulation of cortico-motoneuronal pyramidal output neurons even in those individuals with a caudal precentral hotspot. Even if TMS preferentially targets the rostral part of anterior M1_HAND_ (BA4a) in the posterior crown-lip region, the bulk of cortico-motoneuronal pyramidal output neurons can be expected to be located deeper in the sulcal wall, in the posterior M1_HAND_ (BA4p). We therefore argue that TMS excites the cortico-motoneuronal pyramidal output neurons in the posterior M1_HAND_ (BA4p) always indirectly via an “up-stream” excitation of a neighbouring cortical area regardless of the rostrality of the precentral hotspot. In case of a rostral precentral hotspot location, cortico-motoneuronal pyramidal output neurons are excited via cortico-cortical PMd-to-BA4p projections. In case of a caudal precentral hotspot location, cortico-motoneuronal pyramidal output neurons may additionally excited through excitation of cortico-cortical BA4a-to-BA4p projections.

This model, assuming a different substrate for cortico-cortical excitation depending on the rostrality of hotspot location, makes a testable prediction: It should take longer for the TMS pulse to excite the cortico-motoneuronal pyramidal output neurons, if the precentral hotspot is located more rostrally in the precentral crown-lip region because the primary site of excitation is more “upstream” to BA4p. Our measurements of the MEP latencies showed that this was indeed the case. The more rostral the spatial hotspot location, the longer was the MEP latency, confirming a more “upstream” site of neuronal stimulation in the precentral crown. The results show that “spatial rostrality”, reflected by the rostro-caudal location of the precentral motor hotspot, was paralleled by “temporal rostrality” reflected by a longer MEP latency. This spatial-temporal correlation underscores the functional significance of our TMS mapping results.

The functional relevance of a rostral precentral hotspot location was further corroborated by our task-related functional MRI experiment during which participants performed a visually paced finger tapping task requiring a regular repetition rate at 1 Hz. While the level of task-related activation was comparable for the M1_HAND_, the group with a more rostral hotspot location (i.e., the PMd hotspot group) showed a stronger task-related activation of the PMd than the group with a caudal hotspot location (i.e., M1_HAND_ hotspot group). At the single-subject level, task-related PMd activation showed a positive linear relationship with the spatiotemporal rostrality index. The more rostral the hotspot, the more strongly subjects recruited the PMd during the task. One may argue that the increased PMd activation may be a sign of neuronal inefficiency. In this case, subjects would have to recruit the “inefficient” PMd more strongly to accomplish the task. The behavioural results are at variance with such an interpretation. The relative increase in PMd activity was paralleled by a higher degree of temporal regularity during the visually paced finger tapping task in individuals with a rostral precentral hotspot in the PMd. We therefore argue that the increased PMd recruitment was beneficial, affording a more precise synchronization of the finger tapping with the regular pace provided by the visual cue. Our findings are in good agreement with previous work showing that the PMd plays a prominent role in visuomotor integration (58–63).

Our structural MRI measurements showed that inter-individual variations in hotspot rostrality were also reflected by the microstructural properties of the precentral hand knob. The degree of precentral myelination, indexed by the mean R1-value, scaled positively with the rostrality of the precentral motor hotspot, while hotspot rostrality was not associated with differences in regional cortical thickness. The between-group difference in R1-value was maximal in the anterior lip region of the precentral gyrus. Therefore, it is possible that a higher degree of myelination in the anterior lip region rendered axons in the rostral PMd more sensitive to TMS, shifting the precentral motor hotspot rostrally. Recent structural MRI studies have provided converging evidence that cortical myelination is closely linked to intrinsic functional connectivity (64) and task-related functional activity in unimodal cortical areas, including the visual (65, 66), auditory (67, 68), and sensorimotor cortex (46, 69). This close link between functional properties and regional myelin content provides a good explanation for the structure-function relationship found in this study. A higher precentral myelin content was associated with higher synchronization ability during visually paced finger tapping. Together our results suggest that the degree of regional cortical myelination does not only influence the rostrality of the primary target site of TMS in the crown-lip region of the motor hand knob, but also supports precise temporal integration of visual input and motor output during paced finger tapping.

The results of this study have important implications for the clinical and neuroscientific use of TMS as a non-invasive tool to study corticomotor physiology. Defining the individual precentral motor hotspot location is the standard method for spatial targeting of human M1_HAND_ (6, 17). The prevailing assumption is that hotspot-based, functional targeting secures standardization and thus, reduces inter-subject variability. Here we show that one cannot assume that the same precentral cortical area is targeted when using hot-spot based coil placement. The inter-individual differences in hotspot rostrality introduces gradual differences in the primary target site, resulting in different ratios of primary caudal PMd or rostral M1_HAND_ (BA4a) by TMS. The inter-individual differences in hotspot rostrality also need to be considered in the context of interventional TMS studies in which regular or patterned repetitive TMS protocols are applied to the motor hotspot to induce corticomotor plasticity (70). In recent years, it has been emphasized that the plasticity-induced effects of repetitive TMS targeting M1_HAND_ suffers from substantial inter-individual variability (71, 72). We argue that inter-individual differences in hotspot rostrality may contribute to inter-individual differences in the after-effects on corticomotor excitability. Since inter-individual differences in hotspot rostrality are associated with different structural and functional motor phenotypes, it is possible that individuals with a more rostral or caudal motor hotspot may express different forms of precentral motor plasticity. The influence of hotspot rostrality on TMS-induced corticomotor plasticity warrant further study.

We would like to conclude with some methodological considerations. We only assessed hotspot rostrality in the right non-dominant precentral motor hand knob in right handers. It remains to be shown whether similar findings would emerge, if the dominant motor hand knob was stimulated or if subjects were left-handers. As in our previous sulcus-based mapping studies, TMS mapping used a biphasic rather than a monophasic pulse configuration. Biphasic pulses stimulate cortical neurons more efficiently than monophasic pulses (36). Therefore, we could use lower stimulus intensities which improved the focality of our mapping procedure by using a small-size dedicated coil. Since the second phase of a biphasic TMS pulse dominates the effect on the neural membranes (8, 36), we secured that the second phase of the biphasic pulse induced a posterior-anterior current direction in the precentral gyrus. Future work needs to systematically address how the inter-individual differences in hotspot rostrality are influenced by the orientation of the TMS-induced currents in the precentral hand knob, for instance by using monophasic current configurations. Finally, we would like to stress that the findings regarding hotspot rostrality were found for both TMS target muscles, the FDI and ADM muscle. This consistency provides conceptual within-study replication, which strengthens the reliability of our results. In addition, we identified structural, biophysical, functional and behavioural metrics that scaled with individual variations in hotspot rostrality, providing converging evidence for its functional relevance.

In summary, our multimodal mapping study provides first-time evidence for behaviourally relevant, structural and functional phenotypic variation in the crown of human precentral motor hand knob. We show that the rostrality of the motor hotspot location in the right precentral hand knob scales with structural and functional properties of the precentral gyrus and temporal precision of finger tapping. Our results do not only provide novel insights into the neuroanatomy of the human precentral hand knob, relating brain structure and function to regional excitability and dexterity, they also have important implications for the interpretation of physiological and interventional TMS studies targeting the precentral hand knob.

## Acknowledgments

We thank Sofie Johanna Nilsson for helping with the acquisition of MRI data. We are grateful to Lasse Christiansen, Peter Jagd Sørensen, Silas Haahr Nielsen, Estelle Raffin and Johannes Stelzer for data analysis and comments on the manuscript.

## Author Contributions

R.D. designed the study, acquired, pre-processed, analysed and interpreted the data, and drafted a first version of the manuscript.

K.H.M. designed the study, participated in analysis and interpretation of the data

A.T. designed the study, participated in analysis and interpretation of the data

H.R.S. designed the study, participated in interpretation of the data and in drafting a first version of the manuscript.

---

**Supplementary table 1.** The table lists demographic and electrophysiological data (mean±SEM)

|  | **All participants**<br>**(n=24)** | **M1<sub>HAND</sub> group**<br>**(n=10)** | **PMd group**<br>**(n=14)** | **p-value** |
| --- | --- | --- | --- | --- |
| Sex (Male/Female) | 12/12 | 5/5 | 7/7 |  |
| Age (Years) | 24.2 ± 0.9 | 22.6 ± 1.1 | 25.4 ± 1.3 | 0.11 |
| RMT of FDI muscle (% MSO) | 58.1 ± 2.2 | 55.3 ± 2.4 | 60.1 ± 3.4 | 0.27 |
| **MEP latency at hotspot (ms)** |  |  |  |  |
| Left FDI muscle | 22.6 ± 0.3 | 21.7 ± 0.4 | 23.3 ± 0.4 | 0.007 |
| Left ADM muscle | 22.9 ± 0.3 | 22.0 ± 0.4 | 23.5 ± 0.4 | 0.006 |
| **MEP Amplitude at hotspot (mV)** |  |  |  |  |
| Left FDI muscle | 1.1 ± 0.2 | 1.3 ± 0.3 | 1.0 ± 0.2 | 0.43 |
| Left ADM muscle | 0.5 ± 0.1 | 0.4 ± 0.1 | 0.6 ± 0.1 | 0.5 |
| **MNI-coordinates of motor hotspot** |  |  |  |  |
| Left FDI muscle (x-coordinate) | 38.7 ± 0.6 | 39.4 ± 1.0 | 38.1 ± 0.8 | 0.32 |
| Left FDI muscle (y-coordinate) | -16.6 ± 1.0 | -21.3 ± 1.2 | -13.2 ± 0.6 | <0.001 |
| Left FDI muscle (z-coordinate) | 66.5 ± 0.6 | 65.2 ± 0.9 | 67.4 ± 0.7 | 0.08 |
| Left ADM muscle (x-coordinate) | 35.4 ± 1.2 | 35.8 ± 2.4 | 35.1 ± 1.3 | 0.79 |
| Left ADM muscle (y-coordinate) | -17.8 ± 1.1 | -21.7 ± 1.4 | -15.0 ± 1.0 | 0.001 |
| Left ADM muscle (z-coordinate) | 69.1 ± 1.0 | 67.8 ± 1.6 | 70.1 ± 1.3 | 0.28 |
| **Spatiotemporal rostrality index** |  |  |  |  |
| Left FDI muscle | 0.34 ± 0.05 | 0.14 ± 0.03 | 0.49 ± 0.05 | <0.001 |
| Left ADM muscle | 0.32 ± 0.04 | 0.18 ± 0.03 | 0.42 ± 0.05 | <0.001 |

RMT = Resting motor threshold; %MSO= percentage of maximum stimulation output; MEP = Motor evoked potential; FDI= first dorsal interosseous; ADM= abductor digiti minimi. The p-values refer to between-group comparisons comparing mean values of variables (M1<sub>HAND</sub> vs PMd group).

**Supplementary Figure 1.**
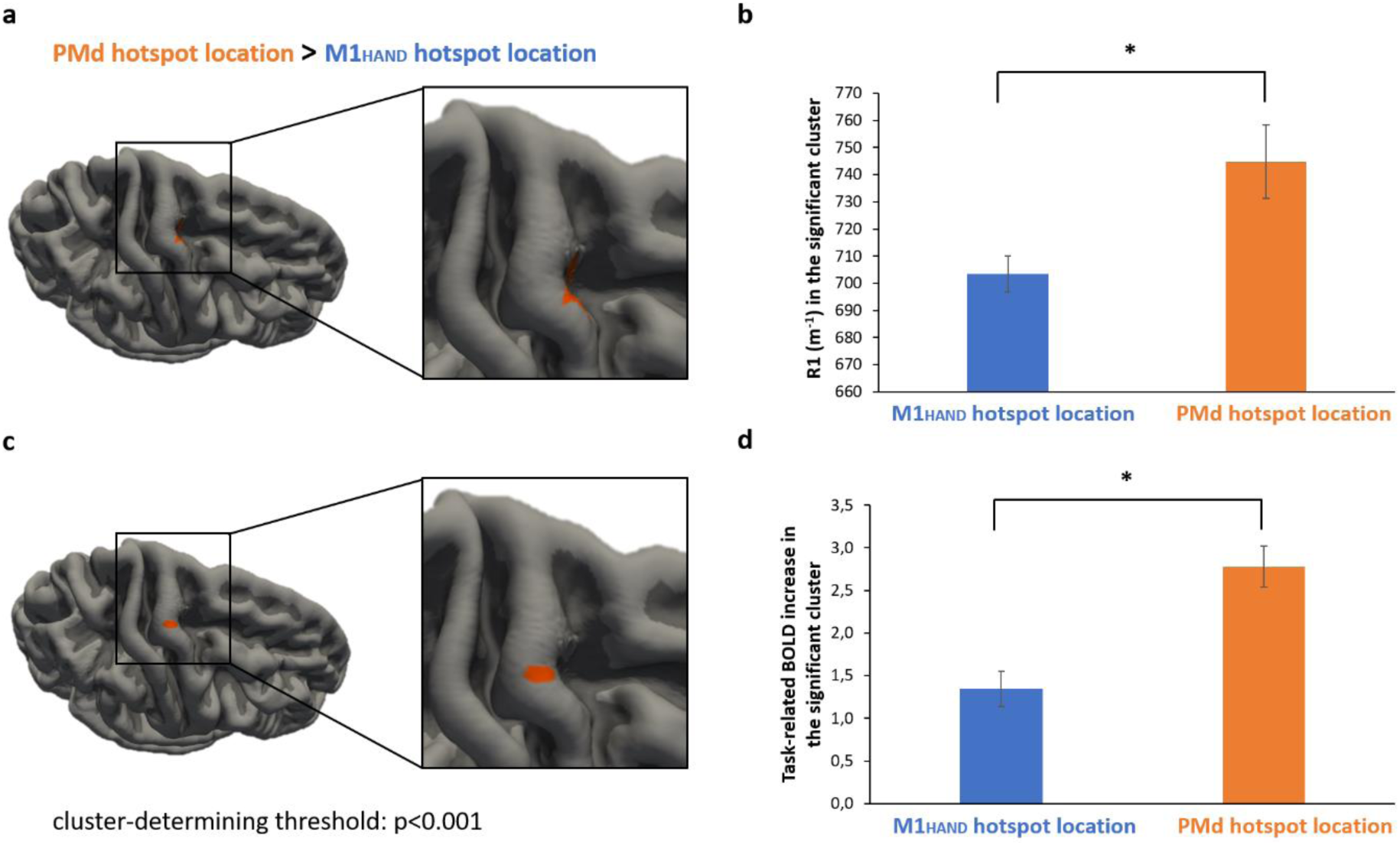
GROUP ANALYSIS OF CORTICAL MYELINATION AND fMRI ACTIVITY IN THE PRECENTRAL HAND KNOB. **a** Surface-based group analysis of R1 estimated cortical myelination indicates that participants with a rostral motor hotspot location have a higher cortical myelination respect to participants with a posterior hotspot location in rostral M1_HAND._ Clusterwise correction for multiple comparisons (cluster-forming threshold of p<0.001) revealed a significant cluster located in the PMd area (cluster wise p< 0.001, coordinates of most significant vertex: x, y, z = 22.2, −12, 58.1). **b** Between-group difference of R1 estimated cortical myelination in the significant cluster. The mean value of R1 is significantly higher in participants with a rostral motor hotspot (orange bar) respect to the subjects with posterior hotspot location in rostral M1_HAND_ (blue bar) (p= 0.022). *= statistically significant. **c** Surface-based group analysis of the mean task-related BOLD increase for both muscles indicates that participants with preferential hotspot location in PMd have a higher functional activation during the visuo-motor synchronization task respect to participants with hotspot location in the M1_HAND_. Clusterwise correction for multiple comparisons (cluster-forming threshold of p<0.001) revealed a significant cluster located on the crown of the precentral gyrus (cluster wise p= 0.0043, coordinates of most significant vertex: x, y, z = 34.4, −15.4, 65.3). **d** Between-group difference of fMRI activity in the significant cluster. The mean value of task-related BOLD increase is significantly higher in participants with preferential hotspot location in PMd (orange bar) respect to the subjects with hotspot location in the M1_HAND_ (blue bar) (p= 0.00018). *= statistically significant.

**Supplementary figure 2.**
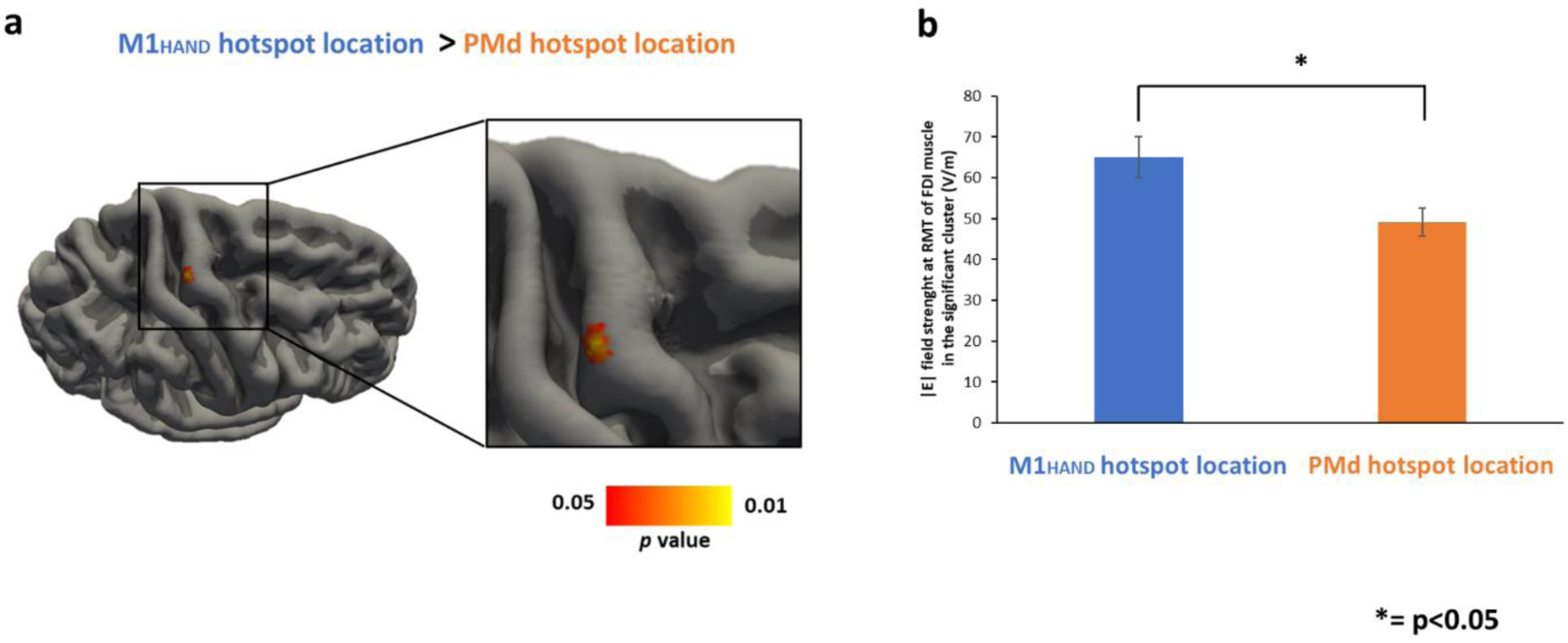
ELECTRIC FIELD SIMULATION IN THE PRECENTRAL GYRUS. **a** Surface-based group analysis of electric field strength |E| indicates that participants with preferential hotspot location in the M1_HAND_ have a higher E-field strength respect to participants with a more rostral location of motor hotpsot. Clusterwise correction for multiple comparisons revealed a significant cluster located mostly in the posterior lip region of precentral gyrus, considering a more liberal cluster forming threshold of p<0,01. **b** Between-groups difference of the electric field strength |E| in the significant cluster shows that the mean value of electric field strength |E| in the subjects with hotspot location in the M1_HAND_ (blue bar) is significantly higher respect to the subjects with preferential hotspot location in the PMd (orange bar), p= 0.019). *= statistically significant.

